# Mapping the malaria parasite drug-able genome using *in vitro* evolution and chemogenomics

**DOI:** 10.1101/139386

**Authors:** Annie N. Cowell, Eva S. Istvan, Amanda K. Lukens, Maria G. Gomez-Lorenzo, Manu Vanaerschot, Tomoyo Sakata-Kato, Erika L. Flannery, Pamela Magistrado, Matthew Abraham, Gregory LaMonte, Roy M. Williams, Virginia Franco, Maria Linares, Ignacio Arriaga, Selina Bopp, Victoria C. Corey, Nina F. Gnädig, Olivia Coburn-Flynn, Christin Reimer, Purva Gupta, James M. Murithi, Olivia Fuchs, Erika Sasaki, Sang W. Kim, Christine Teng, Lawrence T. Wang, Paul Willis, Dionicio Siegel, Olga Tanaseichuk, Yang Zhong, Yingyao Zhou, Sabine Ottilie, Francisco-Javier Gamo, Marcus C.S. Lee, Daniel E. Goldberg, David A. Fidock, Dyann F. Wirth, Elizabeth A. Winzeler

## Abstract

Chemogenetic characterization through *in vitro* evolution combined with whole genome analysis is a powerful tool to discover novel antimalarial drug targets and identify drug resistance genes. Our comprehensive genome analysis of 262 *Plasmodium falciparum* parasites treated with 37 diverse compounds reveals how the parasite evolves to evade the action of small molecule growth inhibitors. This detailed data set revealed 159 gene amplifications and 148 nonsynonymous changes in 83 genes which developed during resistance acquisition. Using a new algorithm, we show that gene amplifications contribute to 1/3 of drug resistance acquisition events. In addition to confirming known multidrug resistance mechanisms, we discovered novel multidrug resistance genes. Furthermore, we identified promising new drug target-inhibitor pairs to advance the malaria elimination campaign, including: thymidylate synthase and a benzoquinazolinone, farnesyltransferase and a pyrimidinedione, and a dipeptidylpeptidase and an arylurea. This deep exploration of the *P. falciparum* resistome and drug-able genome will guide future drug discovery and structural biology efforts, while also advancing our understanding of resistance mechanisms of the deadliest malaria parasite.

**One Sentence Summary:** Whole genome sequencing reveals how *Plasmodium falciparum* evolves resistance to diverse compounds and identifies new antimalarial drug targets.

## Main Text

Malaria has a disproportionately negative impact on human health because its causal protozoan parasites are adept at changing their genomes to evade antimalarial drugs and the human immune system. A human infection may contain upwards of to 10^12^ asexual blood stage parasites. Thus, even with a relatively slow random parasite mutation rate (~10^−9^), within a few cycles of replication each base in the *P. falciparum* genome can acquire a random genetic change that may render at least one parasite resistant to the activity of a small molecule or a human-encoded antibody. The recent evolution of artemisinin-resistant parasites in Southeast Asia now threatens a life-saving treatment in addition to malaria control efforts(1).

Although this rapid evolution impedes our ability to control the disease, it can also be used to our advantage since *in vitro* evolution in the presence of known antimalarials followed by whole-genome sequencing of resistant clones can be used to discover important mediators of drug resistance. For example, the *kelch13* allele that is associated with artemisinin resistance was first identified in the genome sequence of an *in vitro* cultured *P. falciparum* parasite line exposed to sublethal concentrations of dihydroartemisinin in the laboratory(2). This method of resistance evolution can also reveal new antimalarial drug targets(3). Unlike drug targets that are validated using genetic knockdown methods, these chemically validated drug targets are more valuable as it is already clear that their activity can be inhibited in cultured parasites by a small molecule. Furthermore, the inhibitor provides a tool for crystallization and chemical genetic studies. Some of the important new drug targets discovered with this method include lysyl tRNA synthetase (*pfkrs1*)(4), P-type ATPase 4 (*pfatp4*)(5, 6), phosphatidylinositol-4-kinase((7, 8), phenylalanine tRNA synthetase(7), isoleucine tRNA synthetase(9), eukaryotic elongation factor 2(10), and the polyadenylation specificity factor subunit 3(11).

Most studies using this method to date have focused on single gene mutations in response to single compounds even though, in many cases, additional allelic changes have been noted in *P. falciparum* clones during the acquisition of compound resistance. Here, we systematically study patterns of *P. falciparum* genome evolution by analyzing the sequences of clones resistant to diverse compounds with antimalarial activity across the *P. falciparum* lifecycle, in order to comprehensively assess the parasite’s evolutionary development of drug resistance and identify novel antimalarial targets.

## Results

### Next-generation sequencing shows neutral selection for allelic changes in subtelomeric regions and strong positive selection for variants in the core genome

To thoroughly investigate the genomic evolutionary response to treatment with small molecules, we assembled a collection of isogenic *P. falciparum* clones that had acquired resistance to a range of different small-molecule growth inhibitors. Of the 37 different small molecules used during i*n vitro* resistance evolution (Table S1), 26 compounds were hits from *P. falciparum* phenotypic screens(12-15). Others were the result of medicinal chemistry optimization: the spiroindolone NITD678(5), and the imidazolopiperazines GNF179, GNF452, and GNF707(16), which are similar to KAE609 and KAF156, respectively. The latter two compounds are novel antimalarials that are currently in clinical trials((17, 18). Two compounds, atovaquone and primaquine, are licensed antimalarials. Although a few compounds were similar to one another, such as three carbazoles (MMV019017, MMV009063 and MMV665882), and the imidazolopiperazines (GNF707, GNF452 and GNF179), most possessed a distinct variety of functional groups and heterocyclic substructures. Some compounds showed activity across the parasite lifecycle (Table S1, Table S2)).

Although the phenotypes of some clones had been previously described(19), other clones, including those resistant to primaquine and several clones resistant to GNF179, were evolved specificially for this study over a period of 3-6 months (Figure S1) using a slow ramp-up method of compound exposure. The parasite clones exhibited varying levels of resistance relative to their isogenic parent clones (lower and upper quartile of the half maximal effective concentration (EC_50_) fold shift of resistant clones was 2.8 and 37.2, respectively (Table S3)) and in all cases, demonstrated genetically stable resistance. Genomic DNA was available from an average of 6.45 (median = 5) independently-derived clones for each compound.

To identify the genetic basis of drug resistance, 204 clones (including resistant clones and sensitive isogenic parent clones) were resequenced using the paired-end read method of whole genome sequencing. We included 58 published genomic sequences for clones resistant to 12 compounds, including cladosporin(4), atovaquone(20), and GNF179(21). Alignment to the *P. falciparum* reference genome showed that on average, there was 80.3X coverage of the 23.3 million base pair genome with 85.3% of bases covered by 20 or more reads (Table S3). Non-reference alleles present in each evolved group of clones and absent in the isogenic parent clone were identified. In total, we discovered 1,277 single nucleotide variants (SNVs) and 668 small insertions or deletions (indels) that arose in the 262 whole genome sequences during resistance acquisition (Figure1A, Table 1, Table S4).

**Figure 1.**
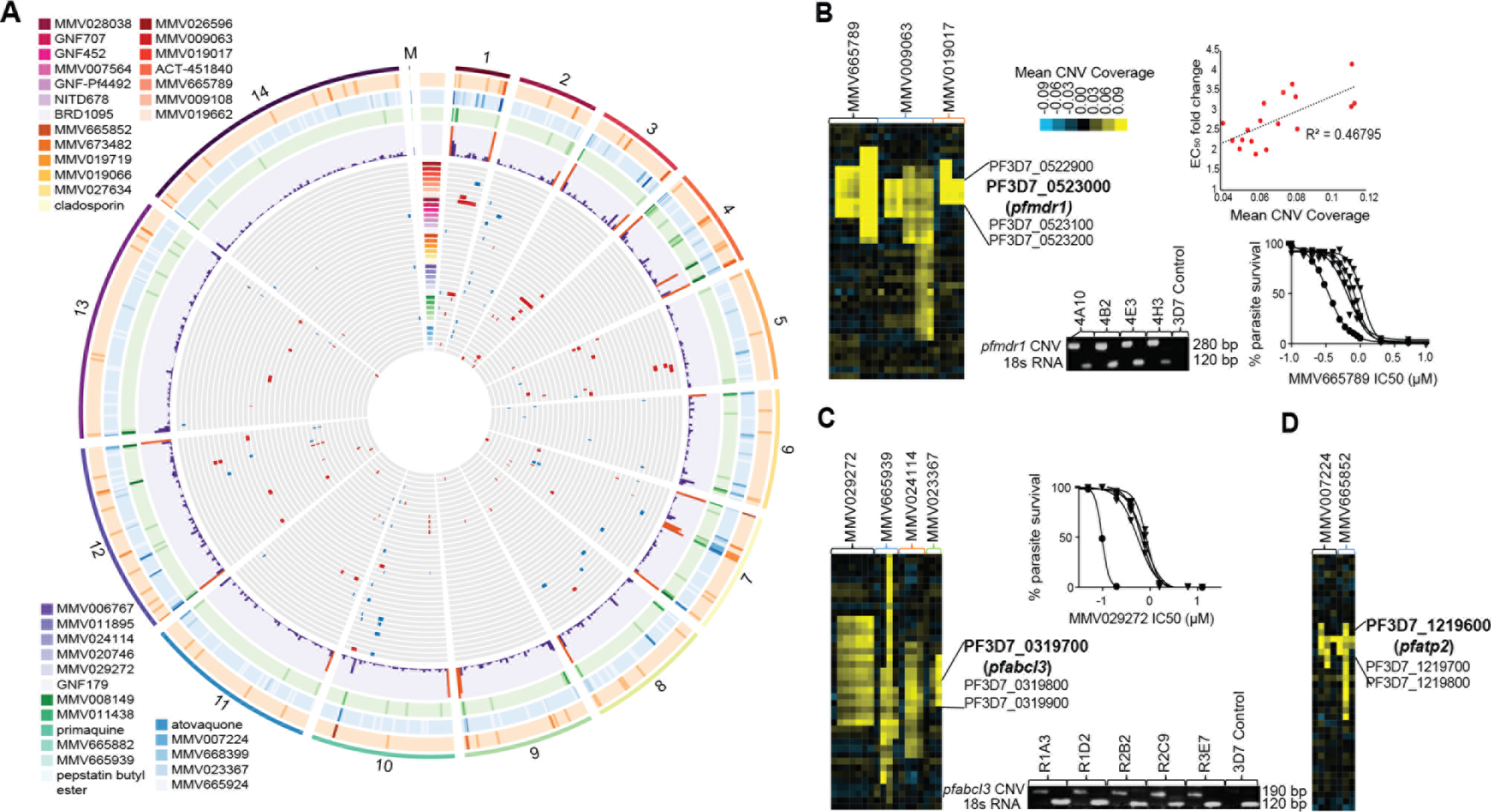
(**A**) Circos plot(68) summarizing single nucleotide variants (SNVs), insertions and deletions (indels), and copy number variants (CNVs) acquired by the 235 *P. falciparum* clones resistant to 37 diverse compounds with antimalarial activity grouped by chromosome. Each bar on the outer 3 rings represents 30,000 base pairs and the darkness of the bar indicates a greater number of mutations. Orange ring: mutations lead to loss-of-function; blue ring: mutations lead to protein modification (non-synonymous change or inframe deletion); green ring: no protein change (synonymous mutations or introns). The purple ring displays a histogram with a count of all three types of mutations, with orange bars if counts exceed 15. The gray rings display variants for resistant clones grouped by compound, with each ring representing one of 37 compounds. Red bars represent the location of CNVs and blue bars represent SNVs. The greatest number of mutations were found in hypervariable genes (*var*, *vir*, and *rifin* families) which are primarily located in the subtelomeric regions (except for additional *var* genes on chromosome 4 and 7). There were notable hotspots on chromosome 1 around the lipid sterol symporter (PF3D7_0107500), on chromosome 3 around *pfabcI3*, and on chromosome 5 at *pfmdr1*. (**B**) CNVs in the known drug resistance gene *pfmdr1*. The heatmap displays clones grouped by compound and is brighter yellow in genes with higher coverage (>2.5X above the standard deviation). The scatter plot shows the correlation between higher CNV coverage and EC50 fold change for clones resistant to all 3 compounds. Resistance shift was demonstrated in MMV665789 resistant clones (diamonds) compared to the sensitive parent (circles). Gel of the PCR products demonstrated that resistant clones contain the *pfmdr1* CNV junction sequence, while a 3D7 control is missing the sequence. (**C**) CNVs in *pfabcI3* are also associated with resistance to multiple compounds. Resistance shift was demonstrated in MMV029272 resistant clones (diamonds) compared to the sensitive parent (circles). Again, the CNV junction sequence was not found in the 3D7 Control DNA compared to clones resistant to MMV029272 (**D**) CNVs were found in *pfatp2* in response to MMV007224 and MMV665882.

**Table 1.** Summary of effects for evolved clones in the core vs non-core regions of the *P. falciparum* genome. Non-core regions were located in subtelomeric regions and internal *var* gene clusters and encoded mostly PfEMP1s, rifins and stevors. Effect changes are as classified using SNPeff(66).

|  | *Effect* | *Number* | *Number* | *Number* |
| --- | --- | --- | --- | --- |
|  |  | **Total** | **Core** | **Non-core** |
| Indels | codon change plus codon insertion | 1 | 1 | 0 |
|  | disruptive inframe deletion | 22 | 17 | 5 |
|  | disruptive inframe insertion | 20 | 11 | 9 |
|  | frameshift variant | 64 | 26 | 38 |
|  | frameshift variant+stop-gained | 1 | 0 | 1 |
|  | inframe deletion | 43 | 37 | 6 |
|  | inframe insertion | 30 | 20 | 10 |
|  | intergenic | 412 | 239 | 145 |
|  | intron variant | 69 | 64 | 5 |
|  | non coding exon variant | 1 | 1 | 0 |
|  | splice region variant+intron variant | 5 | 5 | 0 |
|  | Total Indels | 668 | 449 | 219 |
| SNVs | intergenic | 700 | 178 | 522 |
|  | intron variant | 49 | 30 | 19 |
|  | nonsynonymous coding | 363 | 148 | 215 |
|  | splice region variant+intron variant | 3 | 2 | 1 |
|  | start lost | 1 | 1 | 0 |
|  | stop-gained | 10 | 9 | 1 |
|  | synonymous coding | 1 | 0 | 1 |
|  | synonymous stop | 1 | 0 | 1 |
|  | synonymous variant | 149 | 21 | 128 |
|  | Total SNVs | 1277 | 389 | 888 |
|  | Total Indels and SNVs | 1945 | 838 | 1107 |

Because sequencing might identify false-positive or false-negative alleles, we validated our small variant detection pipeline using several tests. Previous whole-genome analyses of *P. falciparum* using microarrays and sequencing((20, 22) showed that during long-term *in vitro* growth, genes encoding proteins involved in antigenic variation (such as *var* and *rifin* genes), which are usually located in the subtelomeric regions, rapidly acquire mutations. Thus, we hypothesized that we would find a similar pattern for small variants in our data set. Indeed, 1,107 of these variants were located in the 6% of the genome that encodes proteins involved in antigenic variation, referred to as the non-core genome (Table 1). We further predicted that if the mutations in the core genome conferred a selective advantage, then we would find an enrichment of nonsynonymous coding changes. As predicted, the ratio of nonsynonymous to synonymous changes in the noncore region was ~2:1 (216:128) indicating near neutral selection and further confirming the high mutabilility of these genes in the absence of selection. In contrast, the ratio of nonsynonymous coding to synonymous coding alleles in the 23 Mb core genome was 7:1 (148:21) indicating an uneven distribution (p = 7.7E-10) and strong positive selection. In the core genome (containing 838 variants), there were only 148 nonsynonymous changes (Table S5) found in only 83 genes (Table S6), with less than one nonsynonymous change per evolved clone (Table 2).

**Table 2.** Summary of small variants organized by compound used to evolve resistance. Multiple clones from each drug selection were typically sequenced along with at least one isogenic parent clone. In some cases, multiple clones from a single flask were sequenced. These typically yielded virtually identical data, so identical variants from the same round of selection were not counted twice. Core variants are those excluded from subtelomeric regions. Codon changing variants are nonsynonymous coding, splice site, start lost or gained, disruptive insertion or deletion indels and frameshift indels.

| Compound | Variants | Core Var. | Codon changing | No. of clones | Ref/Parent |
| --- | --- | --- | --- | --- | --- |
| ACT-451840 | 19 | 7 | 6 | 4 | 7G8 |
| Atovaquone | 274 | 243 | 21 | 14 | (20) |
| BRD1095 | 8 | 7 | 6 | 7 | Dd2 |
| Cladosporin | 613 | 22 | 5 | 3 | (4) |
| GNF-Pf4492 | 6 | 6 | 5 | 3 | (23) |
| GNF179 | 149 | 87 | 26 | 42 | (40) |
| GNF707 | 30 | 8 | 6 | 3 | Dd2 |
| KAD452 | 8 | 5 | 3 | 1 | (67) |
| MMV006767 | 13 | 12 | 4 | 5 | 3D7 |
| MMV007224 | 25 | 14 | 5 | 3 | 3D7 |
| MMV007564 | 6 | 5 | 3 | 9 | 3D7 |
| MMV008149 | 10 | 8 | 3 | 6 | Dd2 |
| MMV009063 | 24 | 13 | 5 | 8 | 3D7 |
| MMV009108 | 38 | 31 | 3 | 6 | 3D7 |
| MMV011438 | 13 | 4 | 5 | 4 | 3D7 |
| MMV011895 | 58 | 16 | 5 | 4 | 3D7 |
| MMV019017 | 28 | 10 | 6 | 4 | 3D7 |
| MMV019066 | 11 | 11 | 3 | 3 | 3D7 |
| MMV019662 | 92 | 78 | 6 | 15 | 3D7 |
| MMV019719 | 6 | 5 | 3 | 6 | 3D7 |
| MMV020746 | 13 | 13 | 5 | 3 | 3D7 |
| MMV023367 | 6 | 6 | 4 | 5 | Dd2 |
| MMV024114 | 28 | 12 | 2 | 9 | 3D7 |
| MMV026596 | 17 | 12 | 4 | 7 | 3D7 |
| MMV027634 | 19 | 10 | 5 | 3 | 3D7 |
| MMV028038 | 40 | 19 | 6 | 5 | 3D7 |
| MMV029272 | 8 | 8 | 3 | 6 | 3D7 |
| MMV665789 | 7 | 12 | 1 | 4 | 3D7 |
| MMV665852 | 96 | 13 | 5 | 3 | 3D7 |
| MMV665882 | 30 | 23 | 1 | 5 | 3D7 |
| MMV665924 | 6 | 6 | 3 | 3 | 3D7 |
| MMV665939 | 10 | 5 | 2 | 6 | 3D7 |
| MMV668399 | 88 | 45 | 18 | 12 | 3D7 |
| MMV673482 | 8 | 8 | 5 | 4 | Dd2 |
| NITD678 | 72 | 12 | 4 | 3 | (67) |
| PBE | 13 | 11 | 5 | 4 | (39) |
| Primaquine | 53 | 30 | 4 | 7 | Dd2 |
| Parent lines | 0 | 0 | 0 | 27 | NA |
| Total | 1945 | 784 | 206 | 262 |  |

We also predicted that this set of 83 genes would be enriched for genes with annotated roles in drug resistance and that known mutations would be detected in the appropriate compound-specific groups of clones. In fact, gene ontology enrichment testing did reveal that the set of 83 genes was statistically enriched for genes with a *response to drug* gene ontology biological process designation (GO:0042493: p=5.78E-07, Table S7). To the best of our knowledge, there were no false negatives: known variants in P-type cation-transporter ATPase4 (*pfatp4*;PF3D7_1211900)((5, 23) in NITD678-resistant clones, cytochrome bc1 (*pfcytb*;mal_mito_3)(20) in atovaquone-resistant clones, acetyl-coA transporter (*pfact*;PF3D7_1036800) in GNF179-resistant clones, cyclic amine resistance locus (*pfcarl*;PF3D7_0321900)(24) in GNF179, GNF452, and GNF707-resistant clones, and multidrug resistant protein (*pfmdr1*;PF3D7_0523000) in ACT-451840-resistant clones (25) were all detected (Table S6), further validating our data set.

### Most drug-resistant clones acquire copy number variants in addition to SNVs

Copy number variants (CNVs) also contribute to drug resistance in *P. falciparum*(26-28). For example, amplification of *pfmdr1* confers multidrug resistance, including resistance to mefloquine and lumefantrine(29). However, traditional sequencing read depth-based methods of CNV detection are more problematic in *P. falciparum* since the genome is extremely AT-rich (~81%), which leads to uneven genome coverage(30). Thus, we established an automated pipeline for their detection using normalized coverage across our entire data set (see methods). Briefly, average read depth was computed for coding regions only, since intergenic regions of *P. falciparum* are 90-95% AT-rich with AT-repeat segments(31) and thus exhibit reduced alignment confidence. The read coverage data were normalized in three groups depending on the genetic background (3D7, Dd2 or 7G8), allowing the identification of increased read coverage in regions known to be amplified in certain backgrounds (e.g. the *pfmdr1* region amplifications in Dd2). Sets of 2 or more contiguous genes showing ~2.0 X change over the mean were identified. Potential amplified regions were further filtered to yield 159 high-confidence CNVs (core genome, mean average coverage per gene set > ~3.0X over mean coverage and Benjamini-Hochberg corrected p <0.001) (Table 3, Table S8). Altogether, 159 CNVs were observed in *P. falciparum* clones resistant to 27 of the 37 compounds (Table 3, Table S9). CNVs were found primarily on chromosomes 1, 3, 5, and 12 with an average size of ~65 kb (Figure 1, Table 3). Of note, 76 core genome deletions were also identified (Table S10), but most of these were small (< 5 genes), located in subtelomeric regions, and seemed unrelated to the acquisition of drug resistance. For example, some 3D7 clones harbored known deletions of the left arm of chromosome 2, which lacks known drug susceptibility modulators.

**Table 3.**
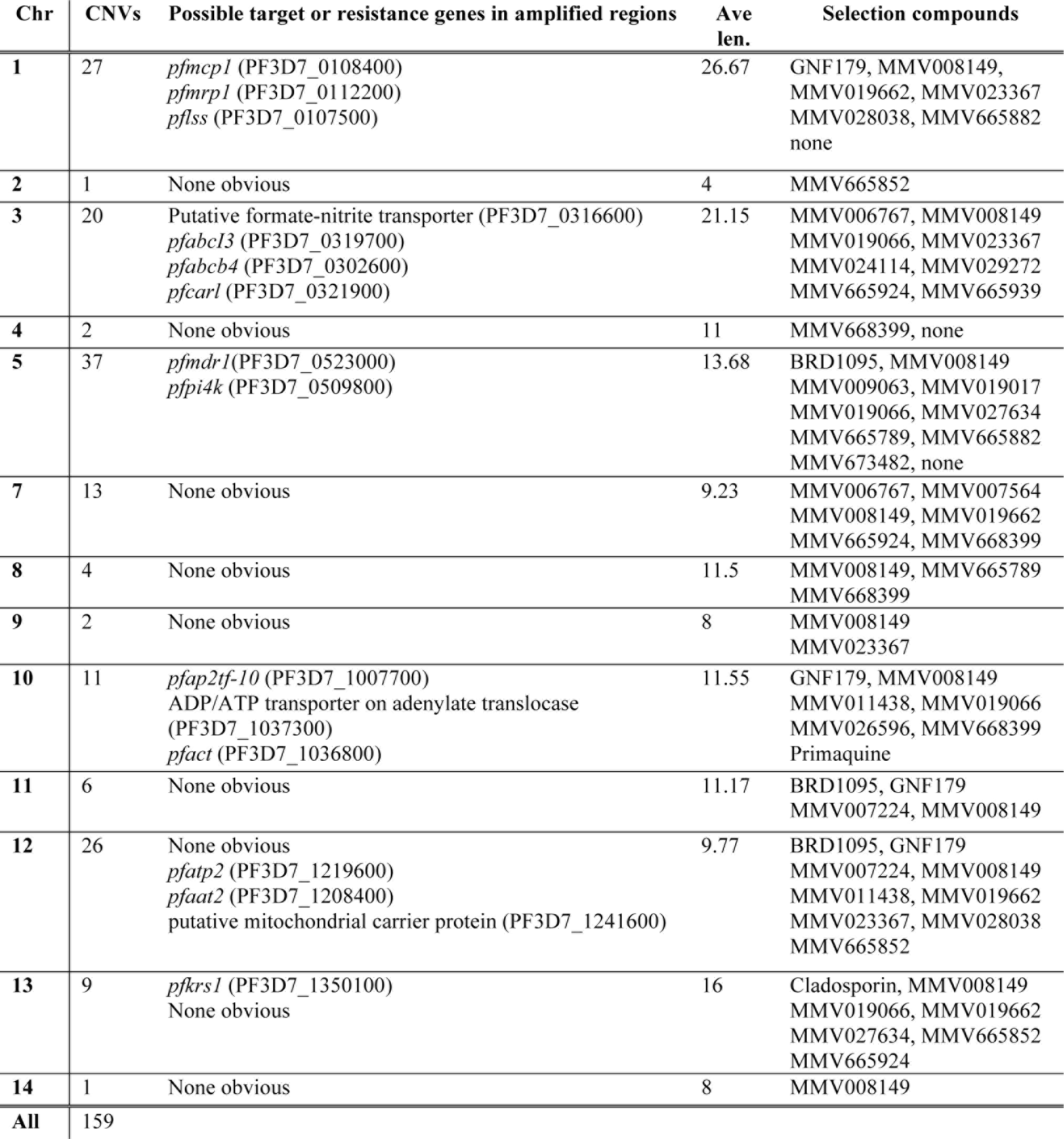
Summary of high quality copy number variants (CNVs) identified in this study. The Ave. Len. is the average size of the amplification defined by the number of genes contained within the region of amplification. Some CNVs covered several plausible candidate genes.

To validate our CNV detection algorithm, we performed additional tests to assess its robustness. We analyzed randomized, permuted average read coverage data and identified only ~8 CNVs fulfilling these criteria. To assess our false negative rate, we next confirmed that we could detect known CNVs previously detected by microarray analysis, including those that span lysyl tRNA synthetase (*pfkrs1*;PF3D7_1350100) in three cladosporin-resistant clones(4) (p = 1.89E-40, 2.29E-23 and 1.28E-12 for the three clones, respectively), and the amplification event encompassing the multidrug resistance protein 1 locus (*pfmrp1*;PF3D7_0112200) in the atovaquone-resistant clone R5a(20)(p = 4.09E-33). The known amplification surrounding *pfatp4* in NITD678-resistant clones(5) was not detected, likely because this clone was sequenced with an older, 60 base pair (bp) read length technology and was therefore not comparable to the rest of the set (although the amplification could be visually detected when compared to other samples sequenced with 60 bp reads). Finally, we used orthologous methods to confirm the CNVs. Many CNVs could be visually confirmed by identifying paired-end reads from sequencing library fragments that spanned the CNV boundaries in the Integrative Genomics Viewer((32, 33) (IGV) and by PCR over the boundaries as previously described (Figure 1B, Figure 1C, Figure S2)(34). We found that CNV read coverage correlated with EC50 shift in resistant clones when no other resistance-conferring mutations were present, such as the *pfmdr1* amplification in clones resistant to compound MMV665789 (Figure 1B).

### Resistance mechanisms are reproducible and structure dependent

As a final test of our dataset we predicted that treating parasites with similar chemical structures would yield reproducible genomic changes. Clustering based on compound similarity did show parasites acquired similar resistance alleles when treated with compounds that are structurally similar (Figure 2). For example, independent clones resistant to imidazolopiperazines (GNF452, GNF707, GNF179) and the closely related compound, MMV07564, acquired mutations resulting in coding changes in the cyclic amine resistance locus (*pfcarl*;PF3D7_0321900). In another example, amplification of the *pfmdr1* region was observed in all 3D7 clones that acquired resistance to three closely related carbazoles: MMV009063, MMV019017, and MMV665882 (Table S8, Figure S2).

**Figure 2.**
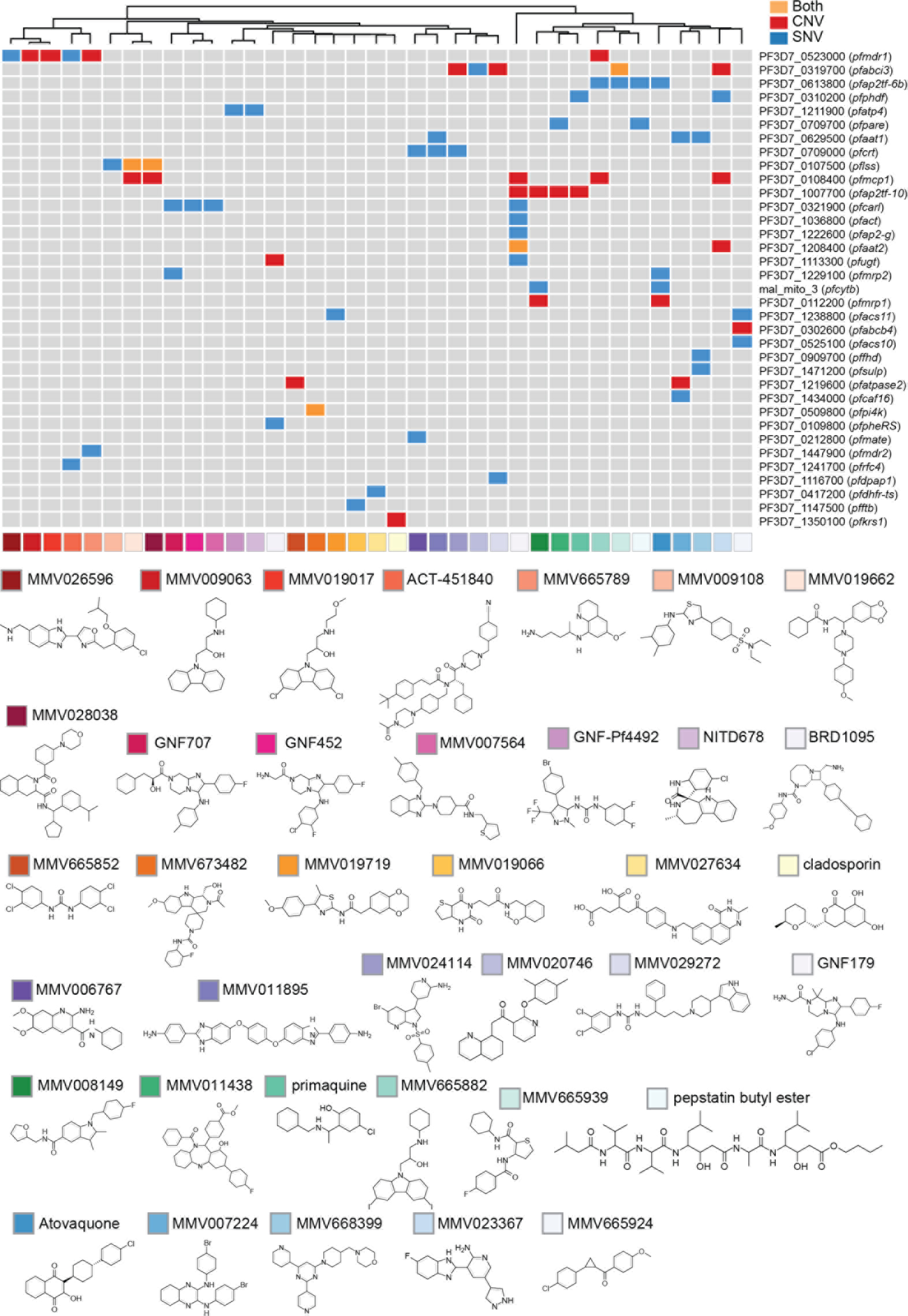
The *P. falciparum* resistome. Compounds were clustered firstly based on their target profile similarities then by their chemical structural similarities. Each compound was assigned a unique color code, which is shared with Figure 1. Genes that were detected as mutated (CNV or SNV or indel) in independently created clones are listed.

### The P. falciparum resistome

Our dataset of CNVs and SNVs revealed 35 genes with two or more types of evidence suggesting a role as a drug resistance determinant or as a drug target (Figure 2). Evidence included: being within an amplification event and bearing SNVs, the presence of two different alleles in the same gene for the same compound, or the same allele appearing in response to two different compounds. Mutations in *pfmdr1* were discovered for 6 different compounds, confirming its known role in drug resistance. Notably, a highly likely resistance mechanism or target gene was discovered for each compound examined. Below, we discuss some of the genes with the strongest evidence.

### Discovery of new alleles in known drug resistance genes

Genetic detection of alleles associated with antimalarial resistance is the primary method of tracking resistance in clinical samples, because *in vitro* phenotypic susceptibility testing is difficult and costly in malaria parasites. Analysis of the mutations present in our large set of resistant *P. falciparum* clones revealed both known and novel genes implicated in drug resistance. In addition to known mutations mentioned above, we found new mutations in several well-known *P. falciparum* resistance mediators. For example, we found four new alleles in the gene encoding the *P. falciparum* chloroquine resistance transporter (*pfcrt*;PF3D7_0709000), including S65R, A138V, K76Q and S90N amino acid changes. Although one of the selection compounds, MMV006767, bears an aminoquinoline core similar to chloroquine, neither MMV011895 or MMV024114 do, providing evidence that PfCRT is a pleotropic transporter regulating drug levels in the digestive vacuole. We also found a novel F806L allele in *pfmdr1* in MMV026596 resistant lines. Furthermore, strong amplification of the *pfmdr1* region was observed in all 3D7 clones that had acquired resistance to MMV009063 (8 of 8 clones, average amplification 10X), MMV019017 (4 of 4 clones) and MMV665882 (5 of 5 clones) as well as MMV665789 (7 of 7 clones) (Table S8). Subsequent testing showed that clones were now cross-resistant to mefloquine, consistent with *pfmdr1* amplification (Figure S3). A new nonsynonymous SNV was also discovered in *pfmdr2* (K840N in response to compound MMV665789). In addition, mutations were detected in gene encoding a plasma membrane-localized ABC transporter (*pfmrp2*;PF3D7_1229100)(35) in a clone that was resistant to imidazolopiperazines (selected with KAD707), as well as in two clones resistant to atovaquone. Changes in *pfmrp2* transcript levels have been associated with levels of *P. falciparum* resistance to quinoline drugs(36), including chloroquine and mefloquine. A SNV was found in the ATP-binding cassette of PfMRP2 at amino acid position D976N in the KAD707- resistant clone, while an A403P change in the transmembrane domain and a frameshift mutation (N1974fs) were detected in the atovaquone resistant lines.

### New drug resistance genes

Knowing the identity of new genes that impart multidrug resistance is important for the design of new drugs, for understanding how existing therapeutics can lose their efficacy in clinical settings, and can help identify likely causative alleles in genome-wide association (GWAS) studies. In our comprehensive data set, we observed that particular genes were mutated repeatedly in response to diverse compounds, indicating that they are more likely to be mediators of general drug resistance. One such candidate is the putative ABC transporter encoded by *pfabcI3* (PF3D7_0319700)(35). Compounds that selected for mutations in *pfabcI3* (MMV020746, MMV023367, MMV024114, MMV029272 and MMV665939) are structurally diverse and some are active across the parasite lifecycle(19). The predicted gene product acquired point mutations (L690I, R2180G, R2180P (two times), and Y2079C) during resistance selections and was central to 12 different chromosome 3 amplification events (Table 2, Figure 1C). All three of the mutated amino acid residues are on loops between two transmembrane domains.

A novel class of potential resistance mediator in this study was a predicted amino acid transporter of the solute carrier family protein (SLC32 family) (pfaat1;PF3D7_0629500). Orthologs of this conserved protein are responsible for the transport of amino acids in synaptic vesicles in humans(37). Mutations in this gene were also found in loops between transmembrane domains and were associated with resistance to three diverse compounds: MMV007224 (-135I inframe insertion), MMV668399 (P380S, K238N, V185L), and MMV011895 ((F230L). This gene was found to be associated with levels of chloroquine resistance in *P. falciparum* in a genome-wide association study(38), and we found that the product of *pfaat1* localizes to the *P. falciparum* digestive vacuole (Figure S4). Thus, the amino transporter encoded by *pfaat1* may play a role in efflux of multiple drugs from the digestive vacuole, similar to PfCRT.

### Nonessential genes and drug activators

Because genes with stop-gained mutations are unlikely to encode critical drug targets, such genes likely play a role in compound detoxification. It was previously reported that the *P. falciparum* Prodrug Activation and Resistance Esterase (*pfpare*;PF3D7_0709700), which is annotated as a lysophospholipase, provides resistance to MMV011438 and pepstatin butyl ester(39). Another gene annotated as a lysophospholipase is PF3D7_0218600, which bears a FabD/lysophospholipase-like domain, and harbors frameshift mutations in two independent clones (MALDA-Primaquine-PQG10 and MALDA-Primaquine-PQA11) that acquired blood stage primaquine tolerance after 5 months of exposure (Figure S1). This gene was not mutated in any of the other 261 clones and comprised two of the eight codon-changing core mutations (3 frameshift, 4 missense, and one stop-gained) in seven different primaquine-resistant clones.

Another amino acid transporter (*pfaat2*;PF3D7_1208400), was mutated in a clone (NMicro-GNF179-S2-3D7-2C) resistant to GNF179, an imidazopiperazine that is closely related to the clinical candidate KAF156(17). *pfaat2* is predicted to contain 10 transmembrane domains and a pfam01490 amino acid transporter domain. The position 903 stop mutation was found in concert with a splice acceptor intronic mutation in the acetyl CoA transporter (*pfact*;PF3D7_1036800), a gene whose disruption confers resistance to imidazolopiperazines(40). CRISPR/Cas9 introduction of the *pfaat2* L903 stop codon into GNF179-sensitive Dd2 parasites revealed that this mutation also conferred resistance to GNF179 on its own (EC50 fold shift of 38- fold compared to the sensitive parent clone) (Figure S5).

### Transcription factors

Several resistance mediators that we discovered are likely to play a role in the parasite’s transcriptional response to drugs. Although there are no examples in *P. falciparum*, work in *S. cerevisiae* and other microbes has shown that mutations in transcription factors can often result in multidrug resistance(41). In our study, the most commonly mutated class of gene was the Apicomplexan AP2 transcription factor family (10 different variants observed in 5 different AP2 transcription factor-encoding genes as well as several potential intergenic promoter mutations). In plants, AP2 transcription factors are involved in the cellular response to stress(42) and in *Plasmodium*, they regulate a variety of developmental transitions including commitment to sexual development and sporogony((43, 44). The most prominent AP2 transcription factor in our set was PF3D7_0613800, which appeared three times with three independent compounds (MMV665882, MMV665939 and MMV011438). In two cases, the same codon deletion (QMEGDNEMEGDNE197Q) was observed, and in one case there was a nonsynonymous codon change (Q197E) at the same position. In addition, a region on chromosome 10 that encompassed a gene encoding another AP2 transcription factor (*pfap2tf-10*;PF3D7_1007700) was amplified in 9 clones (Table 3). The amplification on chromosome 10 appeared in one primaquine-resistant clone (consisting of only 7 genes with an estimated 4 to 5 copies, p *=* 6.6E-5), as well in clones resistant to GNF179, MMV008148 or MMV011438. Variants in the chromosome 6 transcription factor (PF3D7_0613800) were associated with resistance to quinine in genome-wide association studies(45). However, more work is needed to determine whether these changes are associated with multidrug resistance or are related to long term *in vitro* culturing.

The Forkhead-associated domain is a phosphopeptide recognition domain found in some kinases and transcription factors(46). Mutations in an uncharacterized protein bearing this domain (*pffhd*, PF3D7_0909700) were found in all samples resistant to MMV668399 (p = 1E-18 using an accumulative binomial distribution). Mutations included a M1V start-lost mutation, a K392 stop-gained mutation, L95R and G720E missense mutations, and a gene deletion (MALDA-MMV668399-2F2). Given that three of the mutations were predicted loss of function, we hypothesize that this is a gene that inhibits drug action. These same resistant clones also contained mutations in the amino acid transporter (*pfaat1*; PF3D7_0629500) described above, and the inorganic anion transporter (*pfsulp*;PF3D7_1471200) (see below).

### New targets and compound-target inhibitor pairs

In addition to discovering new genes involved in drug resistance, another goal of this study was to identify novel antimalarial drug targets to fill the urgent need for alternative malaria treatments and advance the elimination campaign. In general, we considered mutations in genes encoding enzymes as potential drug target candidates, which was further supported if the mutated gene was compound-specific and docking and homology modeling showed mutations in the catalytic site. A previously reported example is the set of BRD1095-resistant clones, which bear four different amino acid position changes(7) located in the predicted active site of the alpha unit of the cytosolic phenylalanyl-tRNA synthetase.

#### Thymidylate synthase

One likely target-inhibitor pair is dihydrofolate reductase-thymidylate synthase (*pfdhfr-ts*;PF3D7_0417200) and the benzoquinazolinone MMV027634. Although dihydropyrimidine inhibitors of the bifunctional enzyme are well-known antimalarials (e.g. pyrimethamine), all target the dihydrofolate reductase portion of the dual-function protein. We identified three different nonsynonymous mutations mapping to the thymidylate synthase portion of the bifunctional molecule in the MMV027634-resistant lines (Figure 3A). Mapping of the mutations onto a published thymidylate *P. falciparum* crystal structure(47) showed that each mutation flanks the 2’-deoxyuridylic acid (dUMP) binding site of the enzyme (Figure 3A). Independent docking of MMV027634 provided evidence that both substrates occupy the thymidylate synthase active site with a calculated affinity of −8.9kcal/mol.

**Figure 3.**
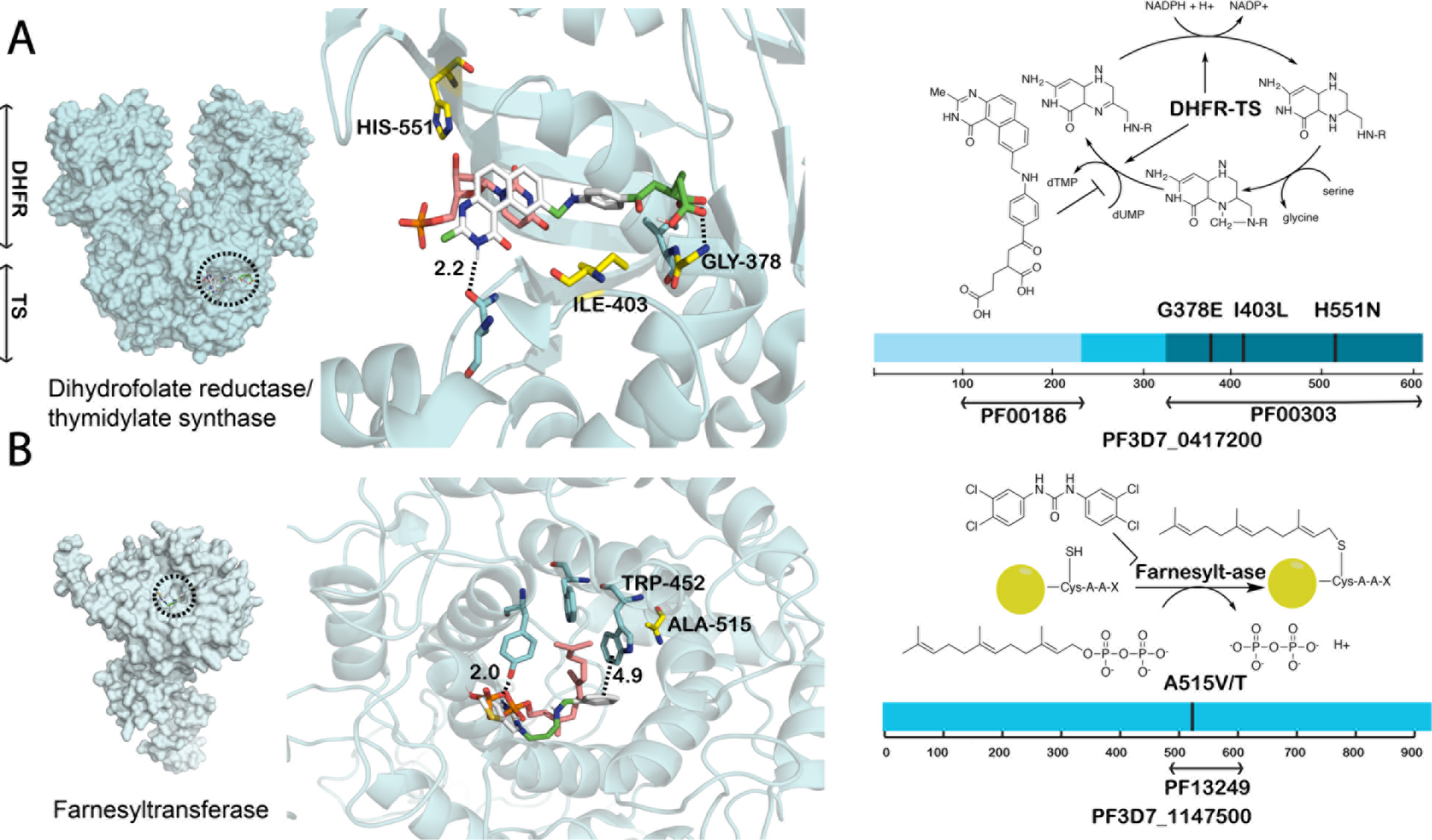
A computational model of active antimalarial small molecules docked against their respective targets. Residues integral to ligand binding are colored cyan, while mutated residues conferring resistance are colored yellow. Schematics of the biochemical pathway, the compound, and conserved pfam domains are shown in the right panel. (**A**) MMV027634 occupies the thymidylate synthase active site. The benzoquinazoline head group of MMV027634 is stabilized by hydrogen-bonding within the dUMP active site of dihydrofolate reductase-thymidylate synthase. Further stabilization occurs at the tail with multiple hydrogen bonds to Gly-378, mutation of which confers resistance to this compound. Mutations are all found in the thymidylate synthase portion of the molecule (http://pfam.xfam.org/family/pf00303). (**B**) MMV019066 and farnesyl pyrophosphate (FPP) are shown concurrently docked within the binding pocket of farnesyltransferase. A model of the *P. falciparum* Ftase beta subunit was constructed using the rat homolog (PDB: 1ZIR) as a template. Despite significant interspecies protein sequence variance, the FPP binding pocket is largely conserved (69). The preferential binding states of FPP and MMV019066 are shown competing for similar hydrophobic space. Resistance mutations are found in the squalene-hopene cyclase domain (http://pfam.xfam.org/family/pf13249).

#### Farnesyltransferase

Farnesyltransferase (*pfftb*; PF3D7_1147500) represents another target gene, which acquired two different mutations in amino acid 515 (A515V, A515T) in three clones resistant to the pyrimidinedione MMV019066 (Figure 3B). Modeling shows that the mutation at amino acid 515 likely disrupts interactions between the thienopyrimidine and the farnesylation active site, resulting in a resistant phenotype. Previous work has shown mutations in the lipid substrate binding site of farnesyltransferase in tetrahydroquinoline-resistant parasites(48).

#### Dipeptidyl aminopeptidase 1

Another potential target gene is dipeptidyl aminopeptidase 1 (DPAP1) (*pfdpap1*;PF3D7_1116700), which encodes an exopeptidase that localizes to the digestive vacuole and cleaves amino terminal dipeptides from proteins or oligopeptides(49). Each parasite clone resistant to MMV029272, an arylurea, carried a mutation (L437S, L415P or N62H) in DPAP1. This gene is considered to be essential as it cannot be disrupted(50). All resistant clones also contained amplifications that encompassed the ABC transporter (*pfabcI3*; PF3D7_0319700), highlighting how resistance mechanisms can be found along with targets (Figure 1C).

#### Aminophospholipid-transporting P-type ATPase

As was previously reported for cladosporin and lysyl tRNA synthase(4), gene amplification events can occur around drug targets. We detected six CNVs in a genomic region on chromosome 12 that encompass a predicted aminophospholipid-transporting P-type ATPase (*pfatpase2*;PF3D7_1219600) (Figure 1D, Table 2), previously named PfATP2. Tandem amplifications with approximate sizes of 8.7 kb, 29 kb, and 25 kb were found in MMV007224-resistant clones and amplifications of 102kb, 95kb and 38kb were found in MMV665852-resistant lines. The 8.7 kb amplification events encompassed only two genes, namely the transporter and a truncated *var* pseudogene (PF3D7_1219500), indicating that the transporter is likely the target. This protein is closely related to PfATP4, an important antimalarial drug target. In the closely related rodent malaria parasite, *P. berghei*, this transporter was refractory to targeted gene deletion attempts, demonstrating its essentiality(51). MMV007224 and MMV665852 are structurally similar to one another and are both active against liver-stage parasites. *S. cerevisiae* orthologs of *pfatp2* encode the Dnf1/Dnf2 aminophospholipid translocases (flippases), which maintain membrane lipid asymmetry at the plasma membrane and contribute to endocytosis (52). In support of this activity, additional nonsynonymous variants in MMV007224-resistant lines were predicted to encode proteins involved in similar processes: Sec24, a component of the coat protein complex II (COPII) that promotes vesicle budding from the endoplasmic reticulum (ER), and Yip1, a protein required for fusion of these ER-derived vesicles with the Golgi. Of note, MMV665852 is a triclocarban, an antibacterial agent whose proposed mechanism is hypothesized to be inhibition of fatty acid synthesis, similar to triclosan, which was proposed to inhibit lipid an membrane function in *Plasmodium*((53, 54).

Finally, in some cases, whether a gene encodes a promising target or a resistance gene may not be obvious. For example, PF3D7_0107500 is annotated as a lipid-sterol symporter in the Resistance-Nodulation-Division (RND) transporter family. We detected 5 different SNVs in this gene in clones resistant to MMV028038, MMV019662 and MMV009108, in addition to 12 amplification events surrounding this gene in clones resistant to MMV028038 and MMV019662. Additional heterozygous SNVs were found in this gene in the MMV019662-resistant clones with chromosome 1 amplification events (Table S11). MMV028038 and MMV019662 both have an amide bond, whereas MMV009108 is very different. RND family transporters are well-characterized in Gram-negative bacteria, where they are involved in extruding toxins, exporting virulence determinants, and maintaining overall homeostasis((55, 56). While specific pumps in this family have been associated with multidrug resistance in bacteria(57), it is currently unclear what specific function the transporter encoded by PF3D7_0107500 has in *P. falciparum*. In addition, three different SNVs (D520Y, K615N, and I596M) were detected in an annotated inorganic anion transporter (*pfsulp*;PF3D7_1471200) in clones resistant to the pyrimidine MMV668399. The probability of randomly observing the same gene three times in 6 selections is extremely low (p = 4.03E-11). PF3D7_1471200 encodes a protein of the *SLC26* membrane protein family(58) with 11 transmembrane domains. Homology models with a fumurate transporter(59) showed that all mutations were located in the STAS domain, a cytoplasmic extension that in some species plays a role in signal transduction.

## Discussion and Conclusions

Our study represents one of the most comprehensive and controlled studies of antimalarial drug resistance acquisition by *P. falciparum* to date. Prior studies examining the parasite’s genetic response during drug resistance development have evaluated the response to known antimalarials only(38, 45, 60, 61), or have focused on coding mutations in one target gene in reponse to a single compound class(4–10, 39). It is likely that the genes that are identified here will prove important to both clinical studies and the process of drug development.

One of the most remarkable findings in this data set is the large enrichment of mutations under positive selection. We focused here on genes for which there were multiple lines of supporting evidence across our study, however it is likely that there are other important genes in the set that were not discussed. Excluding genes contained within CNVs, the list of 57 singleton genes with nonsynonymous mutations includes ones encoding potential drug-able targets, such as GTPases, mRNA decapping enzymes, components of V-type ATPase complexes, kinases, and ubiquitin ligases (Table S5). Although some of these nonsynonymous changes could have arisen by chance, some are plausible. For example, although resistance to ACT-451840 is known to be conferred by mutations in *pfmdr1*(62), a mutation in the gene encoding a putative replication factor C subunit 4 (PF3D7_1241700) that results in a S42T amino acid change near the ATP binding pocket was found in two independent ACT-451840-resistant clones(62) (Figure S6). Replication factor C subunit 4 is the subunit ATPase in the clamp-loading DNA polymerase complex and would be expected to be essential and a good drug target. Modeling shows that ACT-451840 could bind between subunits 1 and 4 of replication factor C, potentially disrupting ATPγ ingress. Our findings also highlight the large number of uncharacterized genes in the *P. falciparum* genome and the need for further functional annotation.

Not all mutations discovered in this study will necessarily confer resistance—they may compensate, specifically or nonspecifically, for a fitness loss caused by a resistance conferring mutation, or they may be associated with adaptation to long-term *in vitro* culture. Genome editing or creating recombinant lines may be used to confirm that alleles provide resistance (5, 7, 8, 11, 25, 40, 63). On the other hand, given the multigenic nature of resistance, it could be difficult to fully recreate a resistance phenotype. For example, although evolved nonsynonymous changes in phenylalanine tRNA synthetase suggested that this was the likely target of BRD1095(7), biochemical assays were ultimately needed to provide proof that this inhibitor blocks protein biosynthesis. We show here that the nonsynonymous changes that emerged in phenylalanine tRNA ligase were accompanied by copy number changes at unrelated sites, potentially explaining why genome editing attempts were unsuccessful.

Although dogma suggests that silent mutations do not affect protein function, the low frequency at which they were found in the set suggested a possible role for some. After excluding genes in the non-core genome, we identified only 21 silent mutations, many of which were in plausible drug targets, including protein kinase 4, Ark3 kinase, cytochrome c oxidase 3, a histone-lysine N-methyltransferase, a putative serine threonine protein kinase, a DNA helicase and an 18S ribosomal RNA. Several of the genes with silent mutations were highly expressed, including cytochrome c oxidase subunit 3 for which the mutation creates a rare codon from a frequently used one (acA (23.8% usage for threonine) to acG (3.6%)). Interestingly, it has been shown that silent mutations in human *mdr1* can confer resistance to cancer drugs by altering protein folding(64). Further work will be needed to establish whether these play a role in drug resistance or adaptation to long-term growth in *P. falciparum*. Similarly, intergenic and intronic mutations were also common. While in most cases resistance was explained by the presence of nonsynonymous coding mutations in the target or in resistance genes, intergenic mutations were also common. Likewise, in our dataset intron variants were much more likely to be found in the core genome than in subtelomeric regions (64:5), suggesting a possible functional role for this abundant class of mutations.

Finally, it is likely that among the genes we identified, several may contribute to clinical resistance at some level. Although the pharmacokinetics are different in humans compared to a tissue culture flask, we repeatedly rediscovered mutations in genes that are known to be important for clinical resistance. It is therefore likely likely that the new genes identified here are prime candidates for genes under selection in clinical isolates. Review of mutations in 3,247 *P. falciparum* clinical isolates from the Malaria Genomic Epidemiology Network *P. falciparum* Community Project reveals that four genes that were identified in this study as mediators of resistance (the ABC transporter *pfabcI3*, the putative amino acid transporter *pfaat2*, the AP2 transcription factor *pfap2tf-6b*, and the farnesyltransferase, *pfftb*), have nonsynonymous to synonymous ratios greater than 2:1, providing evidence that they are likely under positive selection(65)(Table S12). Further research is required to define the potential role of these genes in clinical drug resistance. In addition, our data highlights the importance of CNVs in conferring multidrug resistance and suggests that high-coverage genome sequencing of clinical isolates will provide valuable information on selective pressures in the field.

It is notable that we were able to identify a likely target or resistance gene for every compound that we evolved resistance to using this method. The simple haploid genome and the lack of transcriptional feedback loops suggest that *P. falciparum* is a particularly good model for both target identification and resistome studies. Our characterization of the chemogenetic landscape of *P. falciparum* establishes a new paradigm to guide the design of small molecule inhibitors against deadly eukaryotic pathogens.

## Acknowledgements

This work was supported by grants from the Bill and Melinda Gates Foundation (OPP1040406), the National Institutes of Health (P50GM085764), and the National Institute of Allergy and Infectious Diseases (NIAID) (R01AI103058 and R01AI50234). Sequencing was performed at the Institute for Genomic Medicine Genomic Center, UCSD. Annie Cowell received support from the UCSD Division of Infectious Diseases institutional NIAID training grant (T32AI007036). Erika L. Flannery received support from an NIAID NRSA fellowship (F32AI102567). Victoria Corey received support from a UCSD Genetics Training Program through an institutional training grant from the National Institute of General Medical Sciences (T32GM008666). G.M. Lamonte is supported by an A.P. Giannini Post-Doctoral Fellowship. The red blood cells used in this work were sourced ethically and their research use was in accord with the terms of the informed consents.

## Author’s Contributions

E.A. Winzeler and A.N. Cowell analyzed data, compiled tables and wrote the manuscript. Selections were conducted by G.M. Lamonte, M. Abraham, C. Reimer, V.C. Corey, E.L. Flannery, E. Sasaki, O. Fuchs, A.K. Lukens, E. Istvan, O. Coburn-Flynn, M.C.S. Lee, T. Sakata-Kato, P. Magistrado, C. Teng, S. Bopp, P. Gupta, M. Linares, M.G. Gomez-Lorenzo, C. Cozar, M. Vanaerchot, N.F. Gnadig, J.M. Murithi, and V. Franco. Killing rate assays were performed and analyzed by G.M. Lamonte, M. Abraham, C. Reimer, E.L. Flannery, S.W. Kim, C. Teng, V.C. Corey, E. Sasaki, P. Gupta, E. Istvan, A.K. Lukens, T. Sakata-Kato, P. Magistrado, S. Bopp, M. Linares, C. Cozar, M.C.S. Lee, V. Franco, M.J. Lafuente-Monasterio, S. Prats, and M.G. Gomez-Lorenzo. Circos plot and chemoinformatics analysis was done by O. Tanaseichuk, Y. Zhong, and Y. Zhou. Docking and homology modeling was done by M. Abraham. Sequencing libraries were prepared by S.W. Kim and V. Corey. WGS analysis was performed by A.N. Cowell, V. Corey, and E. L. Flannery. R.M. Williams developed the copy number variant calling pipeline. P. Willis provided compounds. D. Siegel provided information regarding compounds. S. Ottilie coordinated the selection process and exchange of data. L.T. Wang analyzed CNVs by PCR assay. D.E. Goldberg, D.A. Fidock, D.F. Wirth, and E.A. Winzeler supervised the project, provided funding and resources, and helped edit the manuscript. The final manuscript was approved by all authors.

## Data Availability

All relevant sequence data that was not previously published have been deposited in the National Center for Biotechnology Information (NCBI) Sequence Read Archive database with accession code SRP096299. Previously published sequences are deposited in the NCBI SRA with accession codes listed in Table S13.

## List of Supplementary Materials

Table of Contents

Materials and Methods

Figs. S1 to S6

Tables S7, S11, S12, and S13

Additional Data (Captions for Tables S1-S6, S8-S10)

Supplemental References (70-92)

